# Dihydroceramide desaturase promotes the formation of intraluminal vesicles and inhibits autophagy to increase exosome production

**DOI:** 10.1101/2020.11.10.376046

**Authors:** Chen-Yi Wu, Jhih-Gang Jhang, Wan-Syuan Lin, Chih-Wei Lin, Li-An Chu, Ann-Shyn Chiang, Han-Chen Ho, Chih-Chiang Chan, Shu-Yi Huang

## Abstract

Exosomes are important for cell-cell communication. Deficiencies in the human dihydroceramide desaturase gene, *DEGS1*, increase the dihydroceramide-to-ceramide ratio and causes hypomyelinating leukodystrophy. However, the disease mechanism remains unknown. Here, we developed an *in vivo* assay with spatially controlled expression of exosome markers in *Drosophila* eye imaginal discs and showed that the level and activity of the DEGS1 ortholog, *ifc*, correlated with exosome production. Knocking out *ifc* decreased the density of the exosome precursor intraluminal vesicles (ILVs) in the multivesicular endosomes and reduced the number of exosomes released. While *ifc* overexpression and autophagy inhibition both enhanced exosome production, combining the two had no additive effect. Moreover, DEGS1 activity was sufficient to drive ILV formation *in vitro*. Together, DEGS1/Ifc controls the dihydroceramide-to-ceramide ration and enhances exosome secretion by promoting ILV formation and preventing the autophagic degradation of MVEs.

These findings provide a potential cause for the neuropathy associated with DEGS1-deficient mutations.

**Graphical Abstract:** 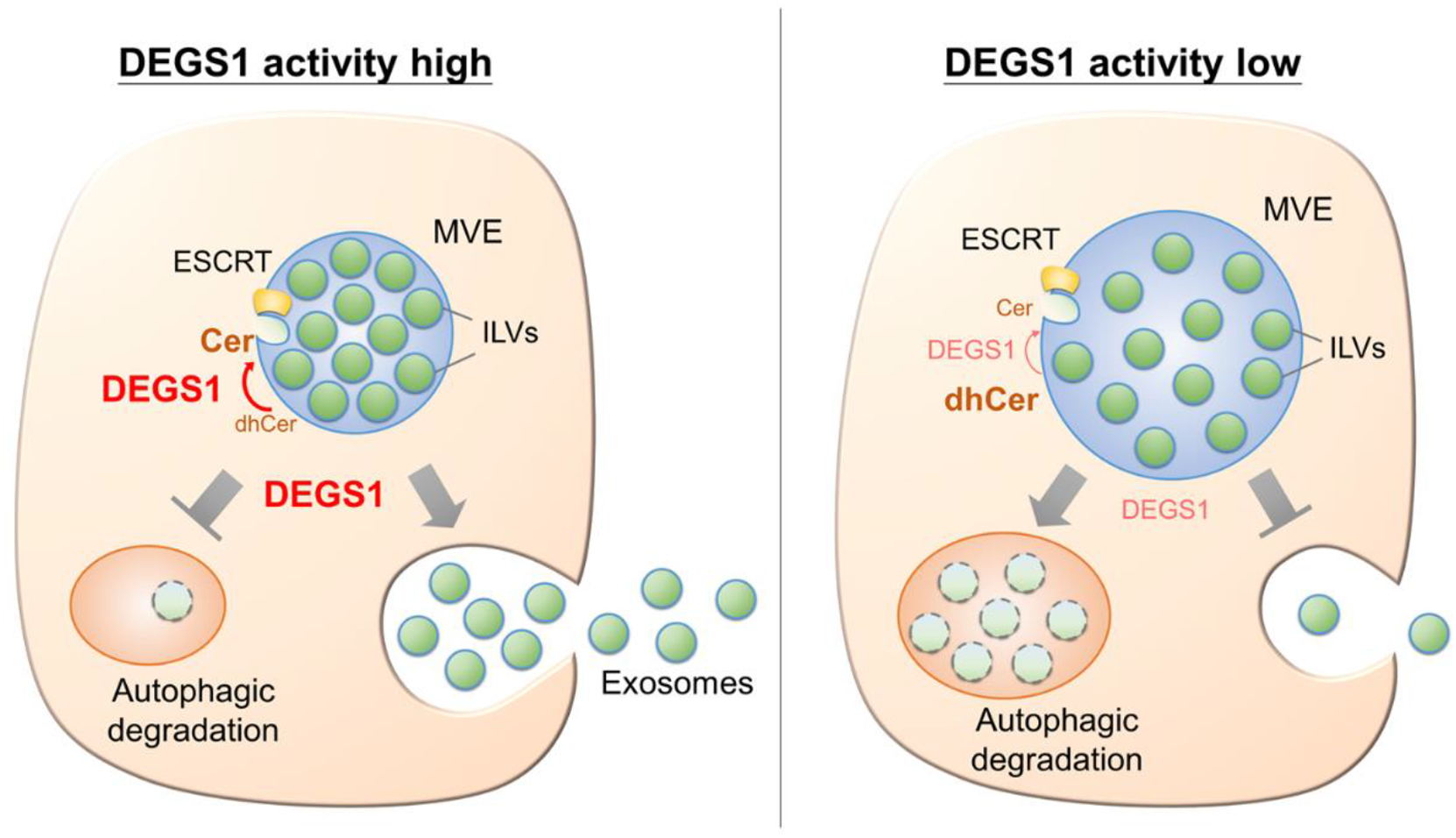

**Highlights:**

- An *in vivo* system was developed for observing exosome production in *Drosophila*.
- Dihydroceramide desaturase (DEGS1/Ifc) promotes exosome production by two means.
- Ifc drives membrane invagination to promote the formation of intraluminal vesicles.
- Ifc inhibits autophagic degradation of MVEs and increases exosome release.

**eTOC Blurb:** The level and activity of dihydroceramide desaturase (human DEGS1 and Drosophila Ifc) correlate with exosome production. Wu et al. show that DEGS1 drives the formation of intraluminal vesicles *in vivo* and *in vitro*. Overexpressing Ifc inhibits autophagy and reduces the degradation of multivesicular endosomes, thus increases exosome release.

## Introduction

Exosomes are secreted vesicles of approximately 100 nm in diameter that mediate cell-cell communication via the transport of proteins, RNAs, and lipid cargos. In the nervous system, exosomes contribute to the formation and maintenance of myelin sheath (Kramer-Albers et al., 2007). Exosomes originate from the invagination of the endosomal membrane in multivesicular endosomes (MVEs) to form intra-luminal vesicles (ILVs) (Raposo and Stoorvogel, 2013). ILVs can form by a mechanism involving the Endosomal Sorting Complexes Required for Transport (ESCRT) complex, which consists of more than 20 proteins that recruit ubiquitinated protein and other cargos and facilitate the budding and scission of the membrane into the MVE lumen to form ILVs (Katzmann et al., 2001, Adell et al., 2014). ILVs can also form via an ESCRT-independent, ceramide (Cer)-dependent mechanism. Trajkovic et al. (2008) show that the inhibition of neutral sphingomyelinase blocks the conversion of sphingomyelin to Cer and reduces exosome release from oligodendrocytes, suggesting that Cer promotes membrane inward budding for ILV formation. When the MVE membrane fuses with the plasma membrane, ILVs are released to the extracellular space as exosomes. Alternatively, ILV-containing MVEs may be targeted for autophagic-lysosomal degradation. Indeed, inhibition of lysosome-autophagosome fusion by bafilomycin A1 has been shown to increases the secretion of α-synuclein in the exosomes (Poehler et al., 2014, Alvarez-Erviti et al., 2011), suggesting cells may use exosomes as a protective mechanism to remove protein aggregates during lysosomal or autophagic dysfunction. Although a ubiquitin-like post-translational modification called ISGylation has been described to promote MVE degradation and thus block exosome secretion (Villarroya-Beltri et al., 2016), little is known regarding the effect of membrane lipid composition on the fate of MVEs.

The final step of Cer *de novo* synthesis is catalyzed by the enzyme dihydroceramide desaturase (DEGS1 in humans and Ifc in *Drosophila*) which inserts a 4,5-*trans*-double bond into the sphingolipid backbone of dihydroceramide (dhCer) to produce Cer. This structural difference has a profound effect on the biophysical properties of Cer; therefore, the relative abundance of Cer versus dhCer may affect the biological properties of cellular membranes (Li et al., 2002). Indeed, genetic or drug inhibition of DEGS1/Ifc has been shown to cause significant accumulation of dhCer which increases reactive oxygen species, induces autophagy and ER stress, causes cell cycle arrest, and even results in cell death in experimental models (Lee et al., 2012). In humans, variants of the human dhCer desaturase gene, DEGS1, have recently been linked to hypomyelinating leukodystrophy and systemic neuropathy (Dolgin et al., 2019, Karsai et al., 2019, Pant et al., 2019). These reports not only confirm an essential role for DEGS1 in the human nervous system, but also show that loss-of-function mutations of DEGS1cause significant accumulation of dhCer and an overall shift of cellular sphingolipid pool towards the dihydro forms (Karsai et al., 2019). However, the underlying cause of neuropathy associated with loss of DEGS1 activity remains to be investigated.

Our previous study shows that eye-specific knock-out of Drosophila *ifc* leads to activity-dependent neurodegeneration, supporting a conserved function in the nervous system. We also show that Ifc can function cell non-autonomously, possibly mediated by exosomes (Jung et al., 2017). Although the cellular level of dhCer is low, several studies report the presence of dhCer in the exosome membrane (Brzozowski et al., 2018, Podbielska et al., 2016, Vallabhaneni et al., 2015). Considering ~30% of cellular Cer is estimated to be produced via the *de novo* synthesis pathway (Lee et al., 2012), whether DEGS1 activity and dhCer level affect exosome formation and MVE degradation to determine exosome production warrants further study.

In the present study, we developed an assay to observe exosome secretion *in vivo* by expressing exosome markers in the dorsal half of *Drosophila* eye imaginal discs. This system allowed the quantification of dorsally made exosomes in response to genetic manipulations on the ventral half of the eye imaginal discs. We also employed the giant unilamellar vesicle (GUV) system to examine if human DEGS1 activity is sufficient to drive the formation of ILVs *in vitro*. Our findings demonstrate an important role for DEGS1/Ifc in regulating both the formation of intraluminal vesicles and the autophagic degradation of MVEs, providing a possible mechanism underlying its function in the nervous system.

## Results

### Knocking out *ifc* reduces the number of exosomes *in vivo*

To evaluate the role of *ifc* in exosome biogenesis, we established an *in vivo* system with the *Drosophila* eye imaginal discs that have been shown to secret exosomes (Gross et al., 2012). We utilized the Dorsal-Eye Gal4 (*DE-GAL4*) expression system (Morrison and Halder, 2010) to drive the expression of the exosome marker GFP-CD63 (Panakova et al., 2005) restricted to the dorsal half of the disc. We were able to detect the presence of GFP-CD63 puncta in the acellular space above the morphogenetic furrow marked by the lack of DAPI staining on the ventral side of the eye imaginal discs (illustrated in Figure 1A), indicating the movement of GFP-CD63+ extracellular vesicles moving along the dorsal-ventral axis. We collected particles released by the discs and determined their sizes by nanoparticle tracking analysis. The average diameter of the secreted vesicles was 113.8 ± 14.3 nm (Figure 1B), consistent with the size range of exosomes. The morphology of the collected vesicles was also consistent with that of exosomes by transmission electron microscopy (TEM) (Figure 1C). Together, these findings show that the labeled puncta exhibit main characteristics of exosomes based on marker proteins, size distribution, and dorsal-ventral movement (Thery et al., 2018), supporting the use of GFP-CD63 as a surrogate to monitor exosome production in the system of *Drosophila* eye imaginal discs

**Figure 1.**
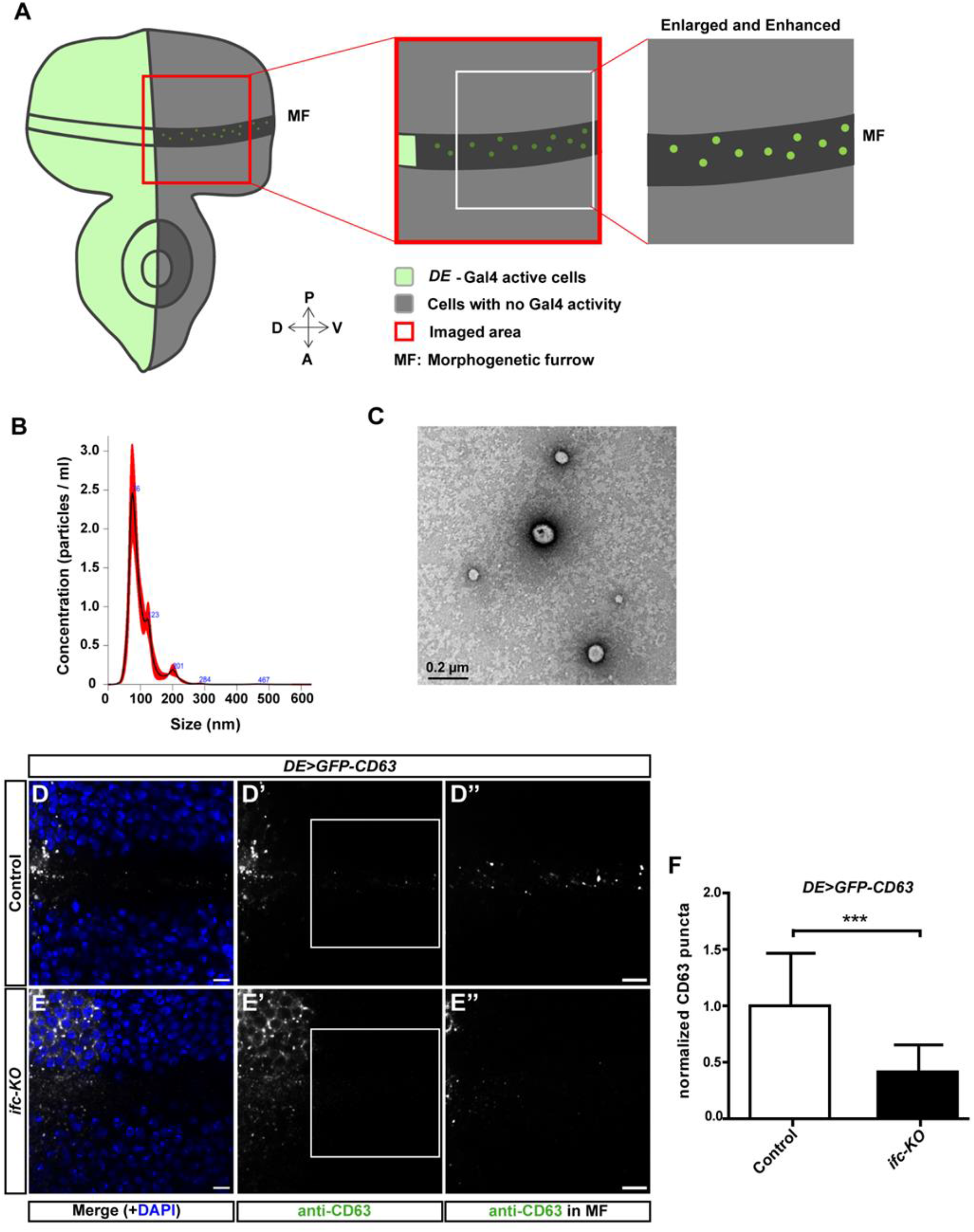
Knocking out *ifc* reduces the number of released exosomes *in vivo*. **(A)** A schematic illustration of the system for observing the release of exosomes. *Dorsal-Eye Gal4* (*DE-Gal4*) drives the expression of *UAS-GFP-CD63* in the dorsal half of the eye imaginal disc (green). The secreted exosomes that move towards the ventral side alone the morphogenetic furrow (MF) into the ventral side where images were taken (red box). The intensity and contrast were linearly enhanced for the quantification of fluorescent puncta in the MF within the region of interest (ROI, white box). **(B)** The representative size distribution of exosomes collected from approximately 50 eye imaginal discs determined by nanoparticle tracking analysis using Nanosight NS300. **(C)** Representative image showing the morphology of exosomes by transmission electron microscopy. **(D-E)** Representative confocal images of *Drosophila* eye imaginal discs with dorsally expressed GFP-CD63 in FRT control (B-B”) and *ifc-KO* (C-C”) showing signals merged with DAPI in blue (B,C), GFP-CD63 detected by anti-CD63 staining in white (B’,C’), and enlarged images of the ROIs with linearly enhanced brightness and contrast to show the GFP-CD63 puncta in the MF (B”,C”). Scale bars: 10 μm. **(F)** Quantification of GFP-CD63 puncta. Bar charts show mean ± SEM of at least three independent experiments. *** p < 0.001 (Student’s *t*-test).

Upon quantification, we found that the number of CD63 puncta detected away from the dorsal compartment was significantly reduced in the *ifc-KO* clones in comparison with the FRT controls (Figures 1D-F). We further utilized this system to express the fusion protein of the flotillin protein Flo2 and RFP as a second exosome marker (Bischoff et al., 2013). Consistent with the results observed with GFP-CD63, *ifc-KO* reduced the number of Flo2-RFP puncta in the ventral compartment (Figure S1). Since *ifc-KO* leads to a drastic accumulation of dhCer, the decrease in exosomes produced by *ifc-KO* clones suggests that the accumulation of dhCer suppresses exosome secretion.

We then examined the morphology of MVE by analyzing the ultrastructural images of MVEs in wild-type (WT) and *ifc-KO* photoreceptors by TEM. The WT MVEs typically contained densely packed ILVs (Figure 2A). In comparison, the MVEs in *ifc-KO* cells appeared dilated (Figure 2B). Indeed, while the numbers of MVE per view field were similar between WT and mutant photoreceptors (Figure 2C, p = 0.3213), the size of the MVEs was significantly larger in *ifc-KO* photoreceptors (Figure 2D, p = 0.0226). Moreover, the average number of ILVs in each MVE was slightly reduced in *ifc-KO*, although the difference from WT was not statistically significant (Figure 2E, p = 0.1153). However, when the size of MVEs was taken into consideration, we found that the density of ILVs was significantly reduced in the mutant MVEs comparing with that of WT (Figure 2F, p < 0.0001). Together, these data show that *ifc-KO* affects the morphology of MVEs.

**Figure 2.**
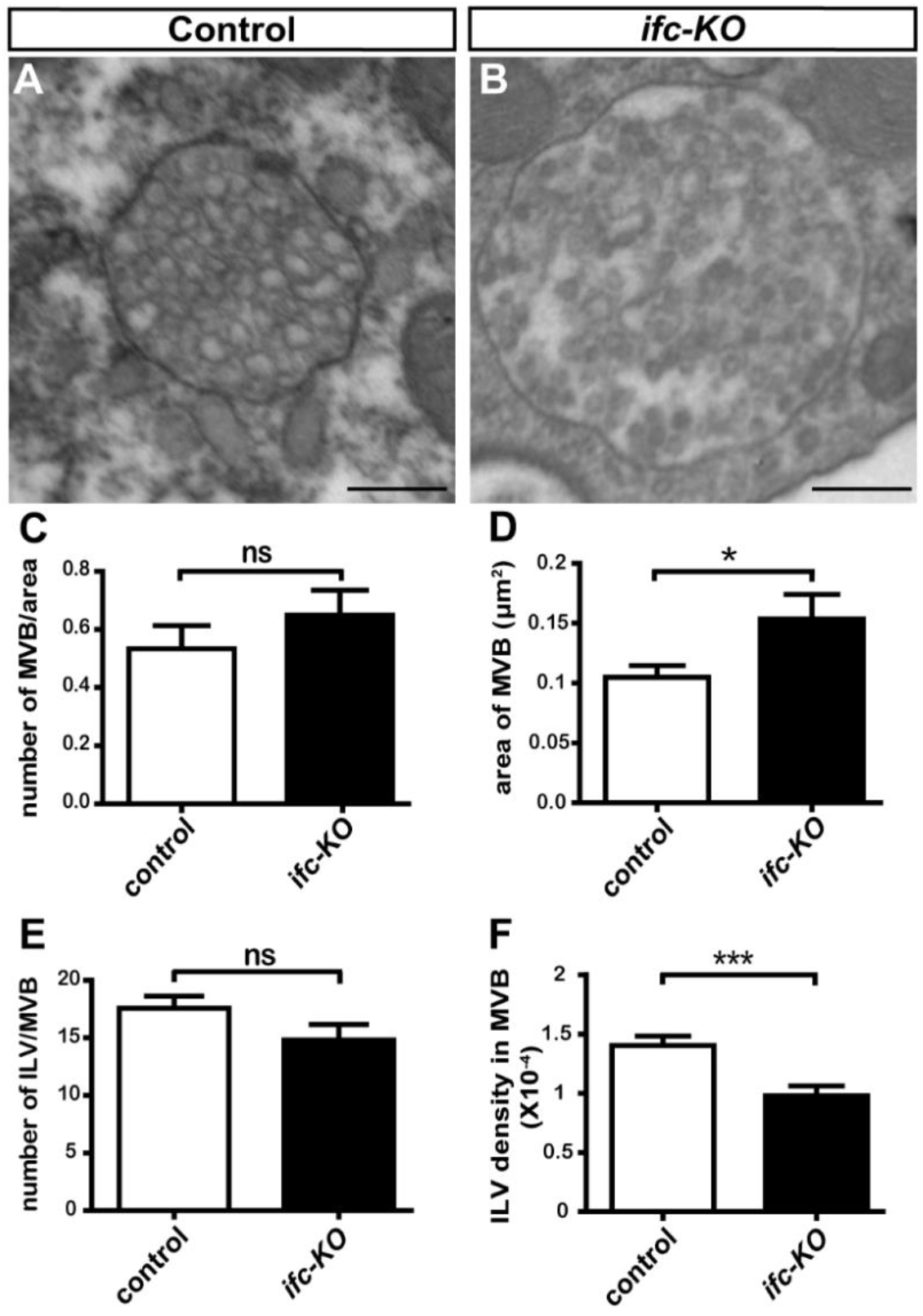
Knocking-out *ifc* reduces the density of intraluminal vesicles (ILVs) inside of multivesicular endosomes (MVEs). **(A and B)** Representative transmission electron microscopy (TEM) images of the wild-type control (A) and *ifc-KO* photoreceptors (B) after 3 days of light stimulation. Scale bars: 250 nm. **(C-F)** Quantification of TEM images including the number of MVEs per field (C), the area of MVEs (D), the number of ILVs per MVE (E), and the number of ILVs per unit area of MVEs (F). Bar charts show mean ± SEM of at least three independent experiments. *** p < 0.001; * p < 0.05; ns: Not significant (Student’s *t*-test).

### Enhancing *ifc* activity increases the number of exosomes

To determine whether enhanced *ifc* activity has the opposite effect of *ifc-KO*, we tested the effect of overexpressing wild-type Ifc with a C-terminal mCherry tag (*ifc(WT)-mCherry*) which we have shown to have the ability to rescue the *ifc-KO* phenotype (Jung et al., 2017). Overexpression of *ifc(WT)-mCherry* increased the number of GFP-CD63 puncta comparing with the UAS control of *mCherry-CAAX* overexpression (Figures 3A,B,D). We performed live imaging of *ex vivo* eye imaginal discs using spinning disc confocal microscopy and light-sheet microscopy and found that, in real-time, *ifc(WT)-mCherry* produced more GFP-CD63 puncta than the *mCherry-CAAX* control (Supplemental videos 1-4). Consistently, overexpression of the *ifc(WT)-mCherry* also increased the number of extracellular puncta labeled with another exosome marker TSG101-HA (Thery et al., 2001) (Figure S2). These results suggest that an increased level of Ifc promotes exosome production.

**Figure 3.**
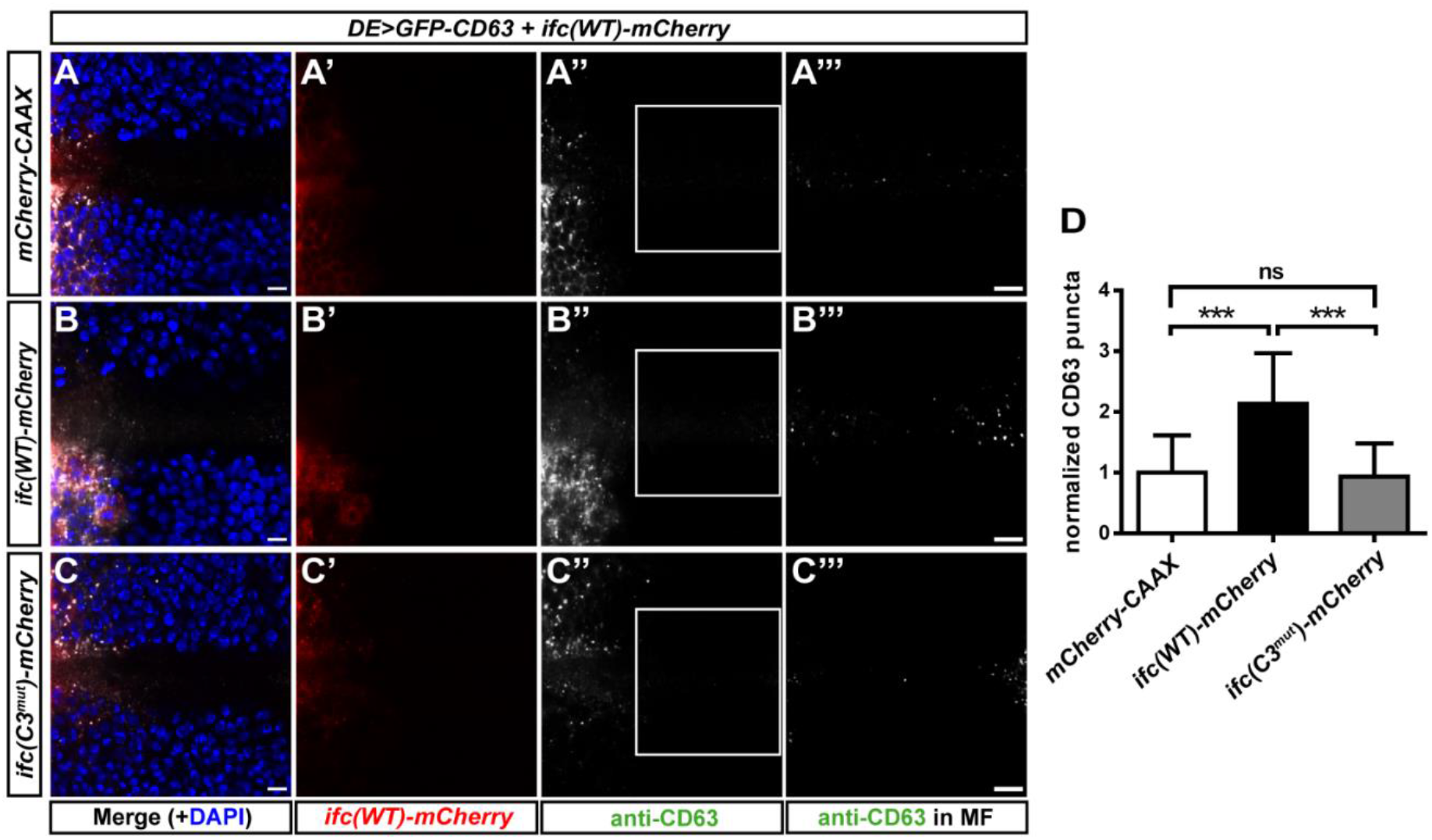
Enhancing *ifc* activity increases the number of exosomes. **(A-B)** Representative confocal images of *Drosophila* eye imaginal discs with dorsally expressed *GFP-CD63* with the UAS control *mCherry-CAAX* (D-D’’’), *ifc(WT)-mCherry* (E-E’’’), or *ifc(C3mut)-mCherry* (F-F’’’) showing merged images with DAPI in blue (D,E,F), mCherry fluorescence in red (D’,E’,F’), anti-CD63 staining in white (D”,E”,F”), and enlarged images of the ROI (white box) with linearly enhanced brightness and contrast to reveal GFP-CD63 puncta in the MF (D’”,E’”,F’”). Scale bars: 10 μm. **(G)** Quantification of GFP-CD63 puncta. Bar charts show mean ± SEM of at least three independent experiments. *** p < 0.001; ns: Not significant (Student’s *t*-test).

To investigate whether the catalytic activity of Ifc was required, we mutated one of the highly conserved histidine boxes that are required for the enzyme activity of Ifc (Figure S3A) (Ternes et al., 2002). The mutation did not affect the protein level as we showed by Western blot that the level of Ifc(C3^mut^)-mCherry was comparable to that of Ifc(WT)-mCherry (Figures S3B,C). Sphingolipidomics result show that *ifc(C3mut)-mCherry* failed to reverse the increase of dhCer-to-Cer ratio in *ifc-KO*, suggesting that the C3 mutation impaired its catalytic activity (Figure S3D). Overexpression of *ifc(C3mut)-mCherry* in *ifc-KO* eye imaginal discs failed to increase the number of GFP-CD63 puncta released to the ventral side in comparison with the *mCherry-CAAX* control (Figures 3A,C,D), suggesting that the desaturase activity of Ifc is required to promote the production of exosomes. These results show that the activity of DEGS1/Ifc to convert dhCer to Cer is required to drive exosome formation *in vivo* and demonstrate a correlation between the level and activity of Ifc with exosome production.

### Ifc interacts with ESCRT genes to regulate exosome production

Since the role of ESCRT proteins in regulating the process of ILV formation is well studied, we investigated the genetic interactions between *ifc* and the ESCRT-0 gene *hrs* and the ESCRT-II gene *vps25*. Because homozygous mutations of *hrsD28* and *vps25A3* are both lethal (Lloyd et al., 2002, Vaccari and Bilder, 2005), we analyzed whether Ifc overexpression genetically interacts with the heterozygous mutations of these two ESCRT genes to regulate exosome production. As expected, the exosomes detected in *hrsD28/+* and *vps25A3/+* were significantly fewer than that in wild-type control (Figure S4). We found that overexpression of *ifc(WT)-mCherry* significantly rescued exosome production in *vps25A3*^/+^ but not in *hrsD28*^/+^ (Figure 4). The result indicates that, for exosome formation, overexpression of *ifc* can compensate for the partial loss of *vps25* but not *hrs*. Since ESCRT-0 (Hrs) is important for the recruitment of ubiquitinated cargos while the ESCRT-II (Vps25) complex is necessary for the initiation of membrane invagination and the recruitment of the ESCRT-III complex (Wollert and Hurley, 2010), this finding supports a role for Ifc in the inward budding of MVE membrane during ILV biogenesis but possibly not in cargo recruitment.

**Figure 4.**
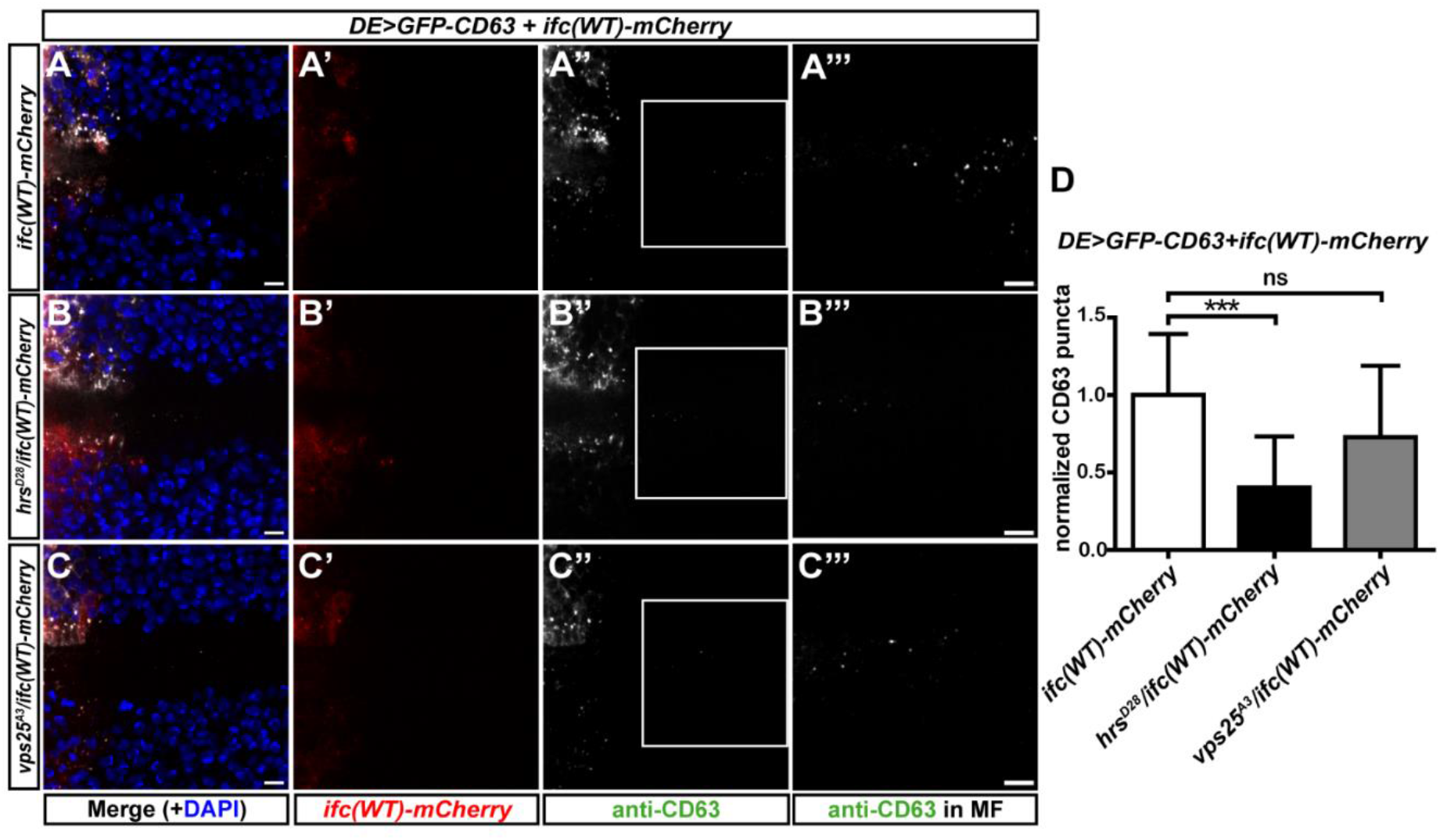
Ifc genetically interacts with ESCRT genes to regulate exosome production. **(A-C)** Confocal images of *Drosophila* eye imaginal discs with dorsally expressed *GFP-CD63* and *ifc(WT)-mCherry* in the control (A), *hrsD28/+* (B), and *vps25A3/+* (C) showing merged images with DAPI in blue (A,B,C), mCherry fluorescence in red (A’,B’,C’), anti-CD63 staining in white (A”,B”,C”), and enlarged images of the ROI (white box) with linearly enhanced brightness and contrast to reveal GFP-CD63 puncta in the MF in the ROIs (A’”,B’”,C’”). Scale bars: 10 μm. **(D)** Quantification of GFP-CD63 puncta. Bar charts show mean ± SEM of at least three independent experiments. *** p < 0.001; ns: Not significant (one-way ANOVA).

### DEGS1 drives the formation of ILVs in GUVs *in vitro*

To evaluate if dhCer desaturase is sufficient to drive ILV formation, we resorted to the *in vitro* model membrane system of GUV. After the generation of GUVs, we tracked the morphology of the GUVs over time and observed no spontaneous ILV formation either with or without the inclusion of dhCer in the lipid mixture (Figures 5A,B-B’,D-D’). Interestingly, when we added human DEGS1 recombinant protein to the GUVs, we were able to observe the formation of ILVs in the GUVs that contained dhCer within 5 minutes (Figures 5A,C-C’,E-E’). When we added the DEGS1 inhibitor fenretinide to the system, the formation of ILVs by DEGS1 was blocked (Figures 5F,J-J’); the ethanol solvent control had no such inhibitory effect (Figures 5F,H-H’). Therefore, the activity of DEGS1 was sufficient to drive the formation of ILV *in vitro* by converting dhCer to Cer. These data further support that the catalytic activity of DEGS1 promotes membrane invagination for the initiation of exosome biogenesis.

**Figure 5.**
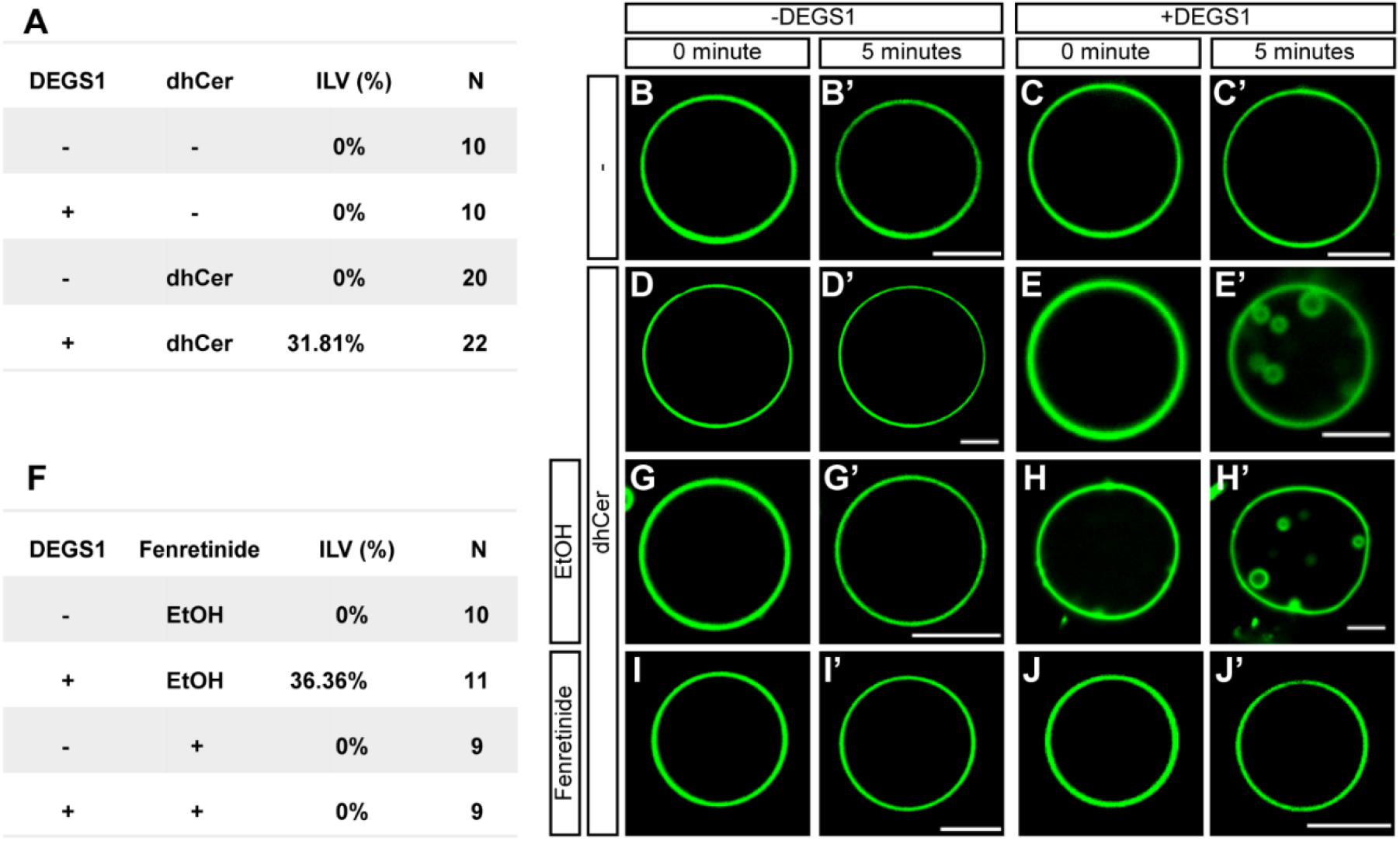
DEGS1 drives the formation of ILVs in the giant unilamellar vesicles (GUVs) *in vitro*. **(A)** Table summary of the percentages of GUVs that formed ILVs within 5 min with or without dhCer and recombinant DEGS1 protein. **(B-E)** Representative images of GUVs in (A). **(F)** Table summary of the percentages of GUVs that showed ILV formation within 5 min with and without DEGS1 protein in the presence of ethanol (EtOH) solvent or 0.2 μM fenretinide, a DEGS1 inhibitor. **(G-J)** Representative images of GUVs in (F). Scale bar: 5 μm.

### Ifc promotes exosome secretion by blocking autophagy-mediated degradation of MVEs

We and others have reported autophagy induction as a common outcome of *DEGS1/ifc* inhibition not only in *Drosophila* but also in mammalian cell lines and animal models (Barbarroja et al., 2015, Holland et al., 2007, Jung et al., 2017, Kraveka et al., 2007). In addition, we previously identified the endosomal localization of Ifc (Jung et al., 2017) and herein showed the regulation of DEGS1/Ifc in exosome secretion. Therefore, we hypothesized that DEGS1/Ifc functions as the pivot of MVB fate between autophagic degradation versus fusion with the plasma membrane for exosome release. To evaluate the interrelationship between autophagy and exosome pathways, we inhibited autophagy by feeding the *Drosophila* larvae 3-methyladenine (3-MA) and chloroquine to block the formation of autophagosome and prevent autophagosomes-lysosome fusion, respectively (Tang et al., 2017, Jung et al., 2017). Inhibition of autophagy by 3-MA and chloroquine both increased the numbers of GFP-CD63+ vesicles released in the wild-type background (Figures 6A-D). Consistent with the results shown in Figure 3, over-expression of *ifc(WT)-mCherry* increased the number of ELVs (Figures 6A,E). Interestingly, feeding 3-MA or chloroquine to larvae overexpressing *ifc* did not further increase the number of GFP-CD63 exosomes in the ventral morphogenetic furrow (Figures 6E-H). These data suggest that GFP-CD63 puncta in MVEs underwent autophagy-mediated degradation in *ifc-KO* and suggest that Ifc promotes the production of exosomes by inhibiting autophagy.

**Figure 6.**
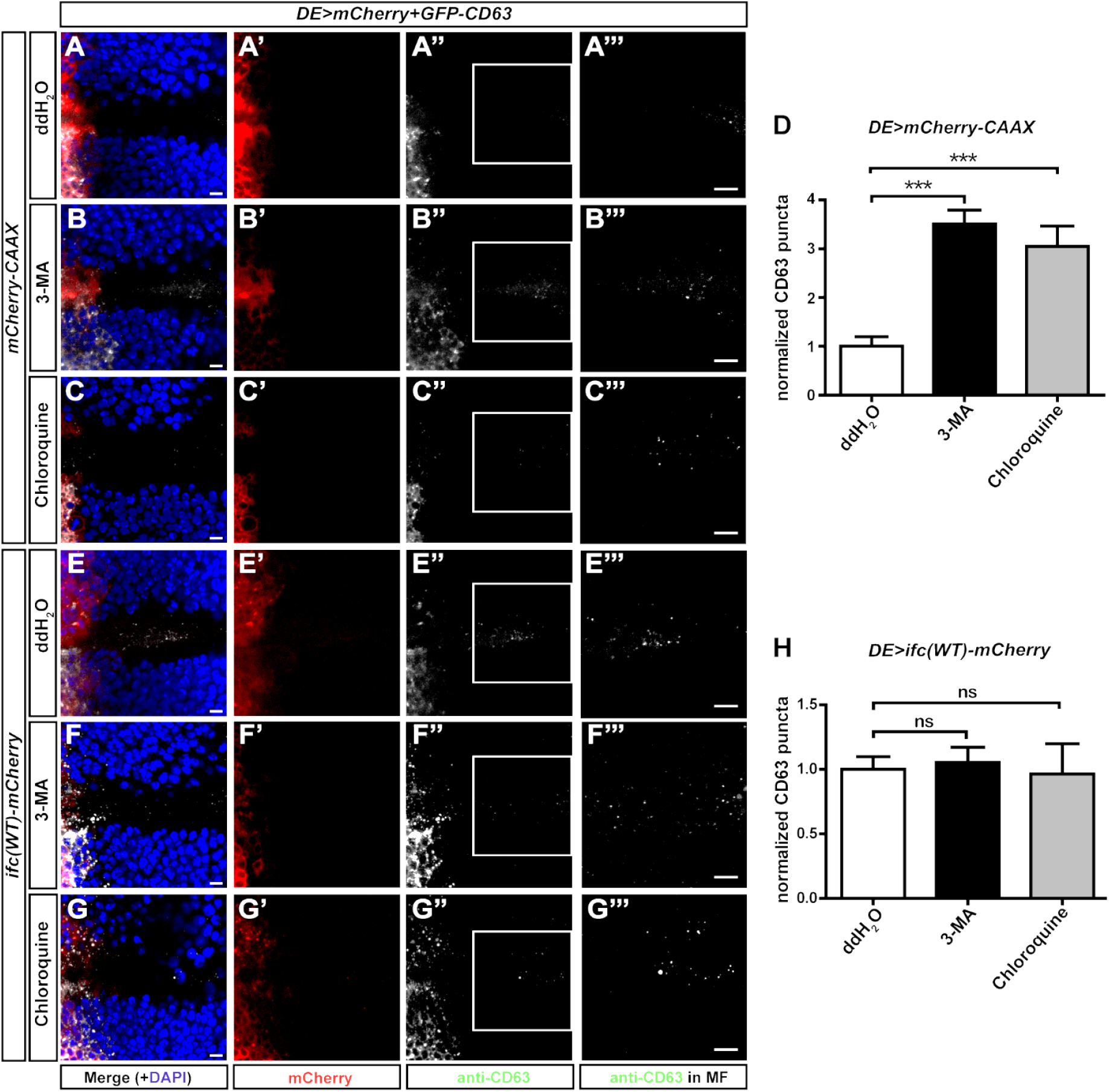
Ifc promotes exosome secretion by blocking autophagy-mediated degradation of MVEs. **(A-C)** Representative confocal images of *Drosophila* eye imaginal discs with dorsally expressed *GFP-CD63* treated with ddH2O (A-A’’’), 3-methyladenine (3-MA, B-B’’’), and chloroquine (C-C’”) showing merged images with DAPI in blue (A,B,C), mCherry in red (A’,B’,C’), anti-CD63 staining in white (A”,B”,C”), and enlarged images of the ROI (white box) with linearly enhanced brightness and contrast to show GFP-CD63 puncta in the MF (A’”,B’”,C’”). **(D)** Quantification of GFP-CD63 puncta. **(E-G)** Representative confocal images of *Drosophila* eye imaginal discs with dorsally expressed GFP-CD63 and *ifc(WT)-mCherry* treated with ddH2O (E-E’’’), 3-methyladenine (3-MA, F-F’’’), and chloroquine (G-G’”) showing merged images with DAPI in blue (E,F,G), Ifc(WT)-mCherry in red (E’,F’,G’), anti-CD63 staining in white (E”,F”,G”), and enlarged images of the ROI with linearly enhanced brightness and contrast to show GFP-CD63 puncta in the MF (E’”,F’”,G’”). **(H)** Quantification of GFP-CD63 puncta. Scale bars: 10 μm. Bar charts show mean ± SEM of at least three independent experiments. *** p < 0.001; ns: Not significant (Student’s *t*-test).

## Discussion

In the present study, we developed an *in vivo* system for the analysis of exosome production using *Drosophila* eye imaginal discs with the dorsal expression of GFP-CD63. The validity of the system was supported by two additional exosome markers, Flo2 and Tsg101, as well as size measurement by nanoparticle tracking analysis. We showed *in vivo* and *in vitro* that the release of exosomes from the dorsal half of *Drosophila* eye imaginal discs to the ventral morphogenetic furrow correlated with not only the level but also the activity of the dihydroceramide desaturase, *ifc*. Our genetic analysis and GUV assay support a clear role for Ifc in the early stage of ILV formation in promoting membrane invagination. In addition, *ifc* also affects the degradation of MVEs via autophagy. Overexpression of *ifc* inhibits autophagy to preserve MVEs thus increase exosome secretion. Together, our data show that DEGS1/Ifc promotes membrane invagination of ILV formation during exosome biogenesis and prevents the autophagic degradation of MVEs, thus promoting exosome formation and release.

Between the ESCRT-dependent and Cer-mediated mechanisms of exosome biogenesis, it is generally believed that the ESCRT-dependent mechanism is involved in the selective sorting of cargos, whereas the Cer-dependent mechanisms are important to control the initial membrane budding (Larios et al., 2020, Teis et al., 2010, Trajkovic et al., 2008). Here, we found that overexpression of *ifc(WT)-mCherry* could rescue the exosome numbers in the heterozygous mutant of *vps25* but not in that of *hrs*, suggesting that *ifc* does not regulate cargo loading. However, *Kajimoto et al. (2013)* show that the continuous Cer catabolism produces sphingosine 1-phosphate that is required for cargo sorting into ILVs in human cells. Exosome membranes share similar lipid composition with lipid rafts, both highly enriched for cholesterol and sphingolipids (Hebbar et al., 2008). Lipid rafts are rigid membrane microdomains that serve as a platform for protein-protein interaction and are essential for protein activation and signal transduction (Lingwood and Simons, 2010). Cer-rich domains have been recognized to affect raft properties by competing and displacing cholesterol (Megha and London, 2004) and promoting the formation of highly ordered gel phases (Bieberich, 2018). The Cer-rich membrane domains may not only serve as a permissive environment to promote membrane invagination for ILV formation, but may also behave as a platform to facilitate the recruitment of ESCRT components. We have previously reported that Ifc predominantly resides in the endosomes in *Drosophila* photoreceptors (Jung et al., 2017), which suggests an on-site conversion of dhCer into Cer for the dynamic regulation of membrane property. How the levels of Cer vs. dhCer on local membranes influence biological processes such as ILV formation need to be further investigated.

Several previous studies have shown that inhibition of autophagosome-lysosome degradation enhances exosome secretion (Danzer et al., 2012, Fader et al., 2008, Iguchi et al., 2016). Interestingly, inhibition of autophagy by Bafilomycin A1 (BafA1) not only increases the secretion of CD63+ exosomes but also increases the detection of autophagic proteins p62 and LC3 in exosomes (Minakaki et al., 2018). However, autophagy proteins ATG12–ATG3 have been shown to interact with the ESCRT-associated protein Alix to promote late endosome to lysosome trafficking as well as exosome biogenesis (Murrow et al., 2015). Also, knockout of ATG5 but not ATG7 significantly reduces the number of exosomes, suggesting that not all of the autophagy core components participate in the regulation of exosome release (Guo et al., 2017). In addition, exosomes purified from cultured cortical neurons treated with BafA1 show a ~2-fold increase in certain sphingolipid species, including Cer and dhCer, indicating BafA1 affects the lipid composition of exosomes (Miranda et al., 2018). Leidal et al. (2020) report that LC3 and LC3-conjugation machinery recruit specific cargos to pack into LC3-positive EVs for secretion, a process that does not require the ESCRT complex but is dependent on the activity of neutral sphingomyelinase 2, an enzyme that hydrolyzes sphingomyelin to produced Cer. In the present study, we showed that exosome production increased with the inhibition of autophagy by either 3-MA or chloroquine. However, combining autophagy inhibition and *ifc* overexpression did not further increase exosome production, implying that *ifc* overexpression inhibits autophagy in opposite to *ifc-KO* which induces autophagy. Membrane Cer/dhCer may play an important role in controlling the fate of MVEs between autophagic degradation and secretion of exosomes; further study is warranted to elucidate the mechanism in detail.

While the effects of Cer have been extensively studied in various contexts, our findings draw attention to dhCer, which is a long-ignored sphingolipid species. Although dhCer is structurally different from Cer only in the lack of the 4,5-double bond, the two sphingolipid species have widely different biophysical properties and bioactivities. Several previous studies have hinted at a role for dhCer in the nervous system. Jung et al. (2017) show that eye-specific *ifc-KO* results in activity-dependent neurodegeneration and demonstrated that *ifc* can function cell non-autonomously. Inherited DEGS1 deficiencies lead to hypomyelination in the central and peripheral nervous systems (Dolgin et al., 2019, Karsai et al., 2019, Pant et al., 2019). Since exosomes are important for myelination and demyelination, whether altered exosome production is the underlying cause of defects seen in patients with inherited DEGS1 deficiencies warrants further investigation.

## Materials and Methods

### Key Resources Table

| REAGENT or RESOURCE | SOURCE | IDENTIFIER |
| --- | --- | --- |
| **Antibodies** |  |  |
| Mouse anti-CD63 | Developmental Studies Hybridoma Bank (DSHB) | Cat# H5C6; RRID: AB_528158 |
| DAPI | ThermoFisher | Cat# D1306; RRID: AB_2629482 |
| Anti-RFP | Rockland | Cat# 600-401-379; RRID: AB_2209751 |
| Anti-HA | Cell Signaling Technology | Cat# 3724; RRID: AB_1549585 |
| Anti-mCherry | GeneTex | Cat# GTX630189 |
| Anti-Actin | Millipore | Cat# MAB1501; RRID: AB_2223041 |
| anti-mouse-HRP | Jackson ImmunoResearch Laboratories, Inc. | Cat# 115-005-003; RRID: AB_2338447 |
| **Drosophila Strains** |  |  |
| ey-FLP | Bloomington stock center (BDSC) | 5622 |
| *cl*, FRT40A |  | 5622 |
| FRT40A |  | 5615 |
| *DE*-Gal4 |  | 29650 |
| UAS-mCherry-CAAX |  | 59021 |
| UAS-hTSG101-HA |  | 77977 |
| *hrs*<sup>D28</sup> |  | 54574 |
| *vps25*<sup>A3</sup> |  | 39633 |
| *ifc*-KO, FRT40A | Jung et al. (2017) |  |
| UAS-*ifc*(*WT*)-mCherry |  |  |
| UAS-EGFP-CD63 | Gift from Dr. Suzanne Eaton<br>(Panakova et al., 2005) |  |
| UAS-*ifc*(*C3*<sup>*mut*</sup>)-mCherry | This study |  |
| UAS-Flo2-RFP | Bischoff et al. (2013) |  |
| **Chemicals and other reagents** |  |  |
| Recombinant human DEGS1<br>protein | Abcam | ab160100 |
| Sucrose | Sigma | S7903 |
| NaCl |  | 31434 |
| Glucose |  | G8270 |
| NADPH |  | 1010782400 |
| ATP |  | A2383 |
| MgCl<sub>2</sub> | Calbiochem | 442611 |

### Experimental Model and Subject Details

#### *Drosophila* strains and genetics

*Drosophila* stocks were maintained at 25°C on standard medium and genetic crosses were performed following standard fly husbandry procedures. Information on individual fly strains can be found on FlyBase (flybase.org) unless otherwise noted in the above Key Resources Table.

### Immunohistochemistry and confocal microscopy

For immunofluorescence staining, larval eye imaginal discs were dissected, fixed, washed, and immunostained with primary and secondary antibodies as previously described (Jung et al., 2017). The antibodies used were listed in the above Key Resources Table. All samples were mounted in Vectashield (Vector Laboratories) and analyzed on a Carl Zeiss LSM880 confocal microscope with LAS AF software. Imaging data were processed and quantified using Photoshop (CS6; Adobe), Illustrator (CS6; Adobe), and ImageJ (National Institutes of Health).

### Nanoparticle tracking analysis

Fifty to 100 eye imaginal discs were dissected and incubated in sterile-filtered Schneider medium (Gibco 21720024) with low-speed agitation (7 rpm) at room temperature for 2.5 h. The supernatant was then separated by low-speed centrifugation at 3,000 x g for 15 min to remove large debris, followed by another centrifugation at 12,000 x g for 15 min. The samples were then passed through a Smart SEC single column (System Biosciences, USA) for EV isolation. The size distribution was determined using NanoSight NS300 (Malvern Panalytical Ltd, United Kingdom) under constant flow (flow rate = 70) at 25°C. The recordings were analyzed by NTA 3.4 software Build 3.4.003 with the detection threshold set to 6.

### Transmission electron microscopy and image analysis of MVE morphology

Exosome samples were prepared as described for nanoparticle tracking analysis. Transmission electron microscopy was performed as previously described (Jung et al., 2017). To characterize the MVE morphology, the fields of imaging for the characterization within the eye of each genotype were selected randomly, and the image files were blinded for the analyses of both the area of MVE and the number of ILV. The area and number of MVE were quantified using Photoshop.

### Timelapse live imaging of exosome secretion

For live imaging of exosome secretion by spinning disc confocal microscopy, timelapse images were aquired using a Zeiss spinning disc microscope equipped with a Yokogawa CSU-X1 scanner unit and a high-resolution CCD (Phtometrics PRISM 95B). Images were acquired through a 40X objective with an exposure time of 200 ms every 5 minutes for 80 min. For live imaging of exosome secretion by light sheet microscopy, the Gaussian intensity distribution of the laser was projecting to an annular ring pattern on the customized aluminum coating mask (thickness of 1,500 angstroms). The masked laser ring image was then projected to a set of galvanometer scanners (6215H, Cambridge Technology) which were composed of a pair of achromatic lenses (Thorlabs, AC254-100-A, Achromat, Ø1”, 400–750 nm), aligned in a 4f arrangement. After passing through the scanning mirror set, the ring pattern was magnified through a relay lens (Thorlabs, AC254-250-A and AC254-350-A Ø1” Achromat, 400–750 nm) and conjugated to the back focal plane of the excitation objective (Nikon CFI Plan Fluorite Objective, 0.30 NA, 3.5 mm WD). The annular pattern was projected to the rear focal plane of the excitation objective and forms a self-reconstructive Bessel beam by optical interference. The signal was detected by a 25X and an sCMOS camera (Hamamatsu, Orca Flash 4.0 v3 sCMOS). Thirty frames were acquired at each time point, with the exposure time of 30 ms for an area of 300 μm x 300 μm. The images for each time point were converted into 4D time series by Imaris 9.1.1. (Bitplane AG).

### Site-directed mutagenesis of *UAS-ifc* mutants

To generate point mutations in the catalytic domain of *UAS-ifc* transgene, the pUASt-ifc was used as a backbone and site-directed mutagenesis was done with QuikChange Kit (Agilent 200523) following the manufacturer’s instruction. All constructs were verified by sequencing. Transgenic flies were generated by embryo injection according to standard protocols of integrase-mediated insertion (Bischof et al., 2007) followed by a germline transmission screen. Primer sequences for *ifc(C3mut)* were: (FWD) GCCAACGAGGCCGCCGACTTTCCG; (REV) GGCGGCCTCGTTGGCGTAGCCCAC.

### Western blotting

The protein extracts were prepared with dissected fly tissues that were homogenized for 10 min at 4°C in 1x RIPA buffer containing protease inhibitors (ThermoFisher A32955), followed by centrifugation at 16,000 x g for 15min at 4°C to remove the debris. The supernatant of the homogenates was mixed with 4X sample buffer (20% glycerol, 4% SDS, 100 mM Tris pH 6.8, 0.002% Bromophenol blue), boiled for 10 min, separated by SDS-PAGE, and transferred in a wet-transfer tank to PVDF membranes (Millipore, IPVH00010) following manufacturer’s instructions (Bio-Rad). PVDF membranes were incubated with 5% non-fat milk in TBST [10 mM Tris (pH 8.0), 150 mM NaCl, 0.1% Tween 20] for 1 h at room temperature and then incubated with primary antibodies at 4°C overnight. Membranes were then washed three times with TBST for 10 min each before incubating with secondary antibodies in TBST for 1 h at room temperature, washed with TBST three times, and developed with ECL reagents (Millipore, WBKLS0500). The signals were captured by the BioSpectrum™ 600 Imaging System (UVP Ltd), and the relative intensities of bands were quantified by densitometry using the Image J software (Schneider et al., 2012).

### Sphingolipidomics

Sphingolipid extraction was performed as described {Jung, 2017 #57}. Briefly, 500 larvae (male and female were equally represented) were snap-frozen in liquid nitrogen and homogenized by grinding.

### Giant Unilamellar Vesicle (GUV) assay

GUV was prepared by the electroformation method (Lee et al., 2018). Briefly, the lipid mixture of Dioleoylphosphatidylcholine (DOPC) and cholesterol was dissolved in chloroform at 1:0.67 molar ratio to the final concentration of 15mM. For lipid mixture with dhCer, the lipid mixture included DOPC:dhCer:cholesterol at 1:1:0.67 molar ratio to the total concentration of 15mM. The lipid mixture was then deposited into a chamber made with a rubber O-ring on an electrically conductive ITO glass. The setup was vacuum dried at 50°C for 1 h, and the solvent was evaporated thereby producing a thin lipid film. The chamber was then re-filled with 2ml of 400mM sucrose. Depending on the experimental conditions, 0.2 μM of the DEGS1 inhibitor fenretinide might be added to the sucrose solution. After sealing the chamber with another piece of ITO glass, the setup was supplied with a sinusoidal voltage waveform of 3.5 V amplitude at 10 Hz frequency and kept at 65°C overnight. The unilamellar vesicles were cooled down and resuspended in 200 μl of 400mM glucose, and confocal images were acquired using Zeiss Cell Observer SD to show the morphology of GUV at time = 0. The recombinant DEGS1 protein at the concentration of 9.62 nM and co-factors (ATP 4 mM, NADPH 2 mM, MgCl2 2.4 mM, NaCl 50 mM, and sucrose 50 mM) were then supplemented to the GUV/sucrose mixture. The post-treatment images were acquired after 5 min of incubation at 37°C.

### Quantification and statistical analysis

Quantitative data were analyzed using a two-tailed unpaired Student’s *t*-test or one-way ANOVA with Tukey’s multiple comparison test. Data analysis was performed using GraphPad Prism 5. All data in bar graphs are shown as mean ± SEM. The quantified GFP-CD63 puncta in the experimental groups were normalized to the average of the controls which was set as 1. *P* values of less than 0.05 were regarded as statistically significant. All quantifications included at least three experimental replicates.

## Acknowledgments

The authors are grateful to Drs. Guang-Chao Chen, Robin Hiesinger, Jennifer Jin, Guang Lin, Ya-Wen Liu, Yi Henry Sun, Chao-Wen Wang, Chi-Kuang Yao, and Mr. Fei-Yang Tzou for their comments on the manuscript. The authors sincerely thank Dr. Ya-Wen Liu and Ms. Yu-Chen Chang for helping us established the GUV system. We thank the staff of the Imaging Core at the First Core Lab, National Taiwan University College of Medicine, the Second Core Lab, Department of Medical Research, National Taiwan University Hospital, and the Electron Microscopy Laboratory, Tzu Chi University, for technical support. We thank WellGenetics, Inc. for embryo injection service. Stocks obtained from the Bloomington Drosophila Stock Center (NIH P40OD018537) were used in this study. This work was supported by the Ministry of Science and Technology of Taiwan grants 107-2635-B-002-001 to S.-Y. Huang and 108-2311-B-002-011-MY3 to C.-C. Chan, National Taiwan University Hospital grant 109-21 to S.-Y. Huang, and National Taiwan University grant 109L104308 to C.-C. Chan.

## Author contributions

S.-Y. Huang and C.-C. Chan conceived the study; C.-Y. Wu, J.-G. Jhang, W.-S. Lin, and C.-W. Lin carried out the fly experiments; J.-G. Jhang developed the *in vivo* exosome release assay; C.-Y. Wu and H.-C. Ho carried out the EM analysis; C.-Y. Wu, L.-A. Chu, and A.-S. Chiang performed live imaging; C.-Y. Wu designed and carried out the GUV experiments; C.-Y. Wu, W.-S. Lin, and S.-Y. Huang prepared the figures; S.-Y. Huang wrote the first draft; S.-Y. Huang, and C.-C. Chan reviewed and edited the manuscript with comments from all other authors; S.-Y. Huang and C.-C. Chan obtained the funding for the project; S.-Y. Huang and C.-C. Chan supervised the project.

## Declaration of interests

The authors declare no competing interests.

## Abbreviations

Cer: ceramide
dhCer: dihydroceramide
ESCRT: Endosomal Sorting Complexes Required for Transport
GUV: giant unilamellar vesicles
ILVs: intraluminal vesicles
MVEs: multivesicular endosomes

---

**Figure S1.**
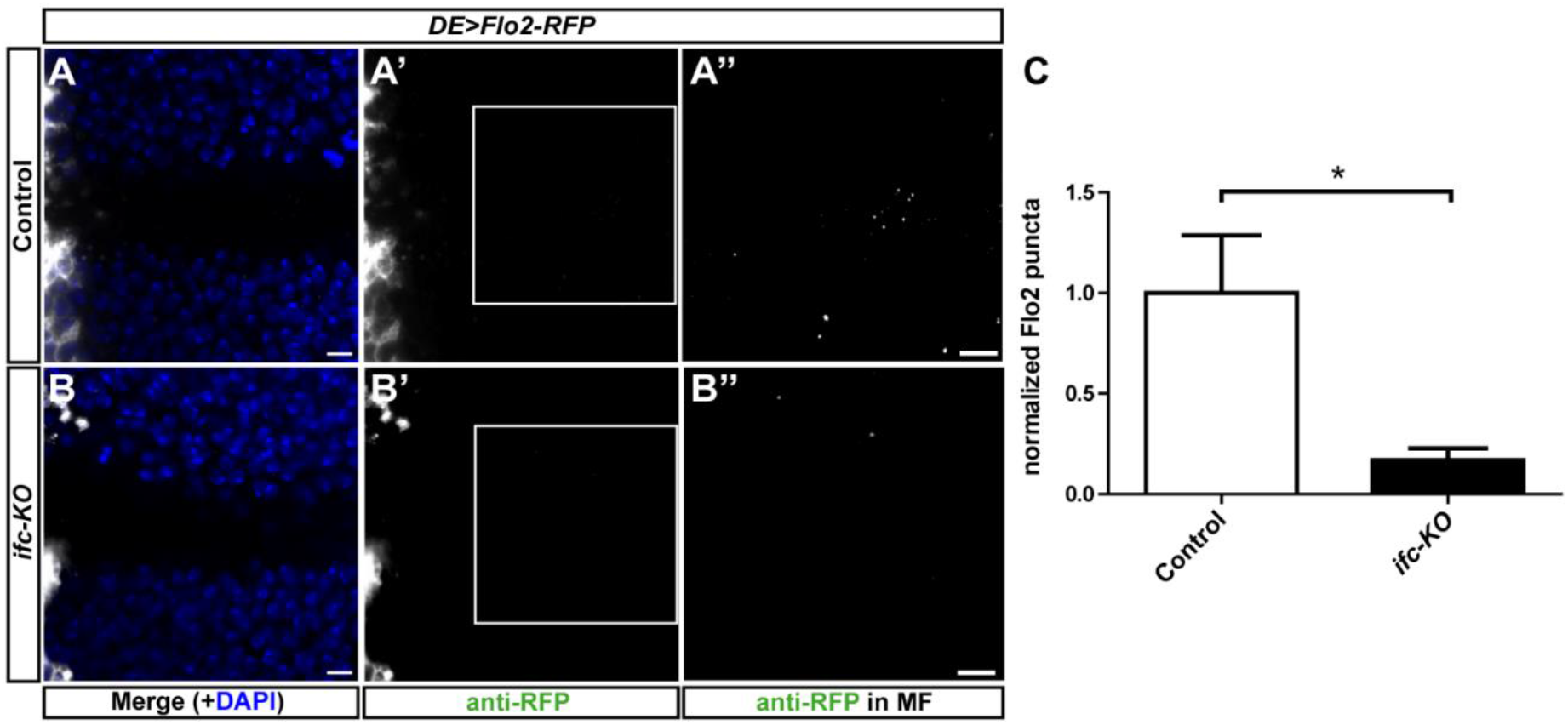
Knocking out *ifc* reduces the number of Flo2-labelled exosomes. **(A-B)** Representative confocal images of *Drosophila* eye imaginal discs expressing Flo2-RFP in the dorsal compartment in the FRT control (A-A’’) and *ifc-KO* eyes (B-B’’) showing merged images with DAPI in blue (A,B), Flo2-RPF signal enhanced by anti-RFP staining pseudocolored in white (A’,B’) and enlarged and linearly enhanced images of the ROI (white box) to reveal Flo2-RFP puncta in the MF (A”-B”). Scale bars: 10 μm. **(C)** Quantification of Flo2-RFP puncta. Bar charts show mean ± SEM of at least three independent experiments. * p < 0.05 (Student’s *t*-test).

**Figure S2.**
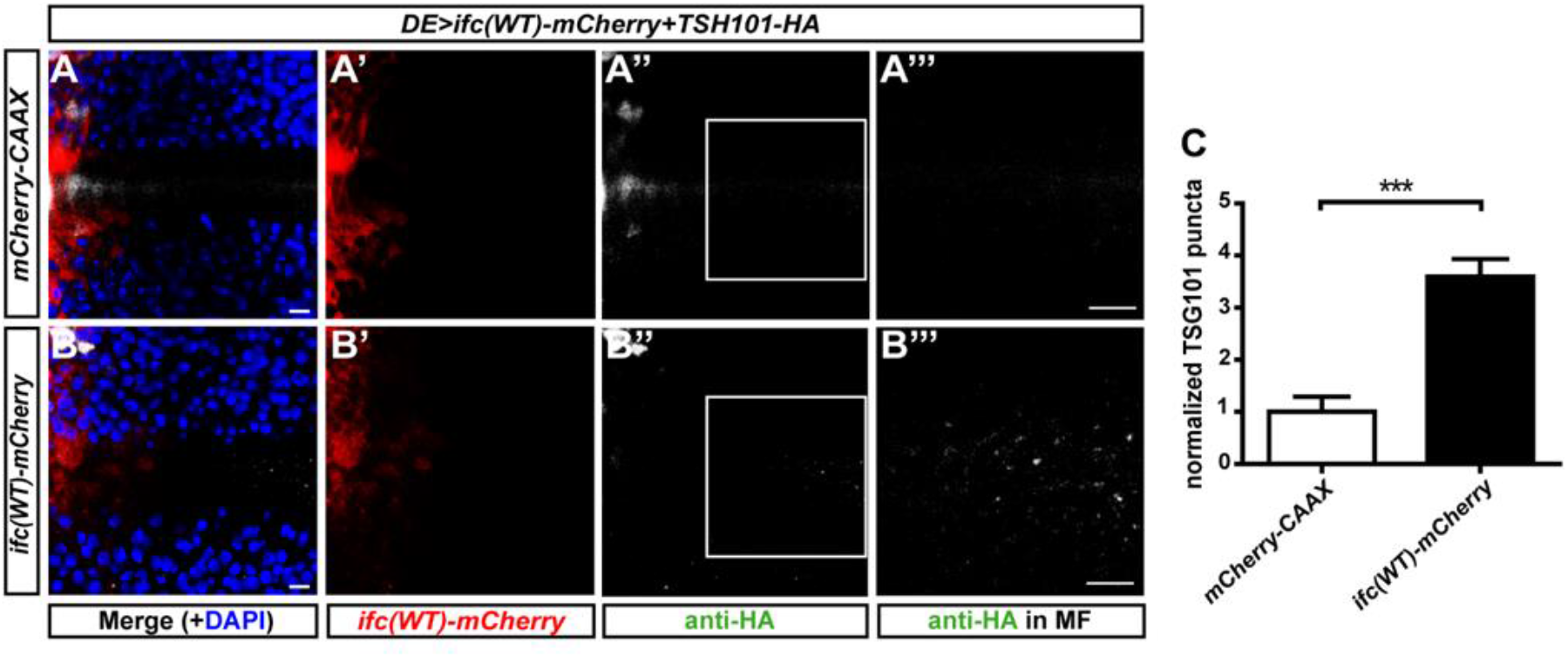
Enhancing *ifc* activity increases the release of TSG101-HA-labelled exosomes. Representative confocal images of *Drosophila* eye imaginal discs dorsally expressed *TSG101-HA* with UAS *mCherry-CAAX* (control, A-A’’’) or *ifc(WT)-mCherry* (B-B’’’) showing merged images with DAPI in blue (A,B), mCherry fluorescence in red (A’,B’), anti-HA staining in white (A”,B”), and enlarged images of the ROI (white box) with linearly enhanced brightness and contrast to reveal TSG101-HA puncta in the MF (A’”,B’”). Scale bars: 10 μm. **(C)** Quantification of TSG101-HA puncta. Bar charts show mean ± SEM of at least three independent experiments. *** p < 0.001; ns: Not significant (Student’s *t*-test).

**Figure S3.**
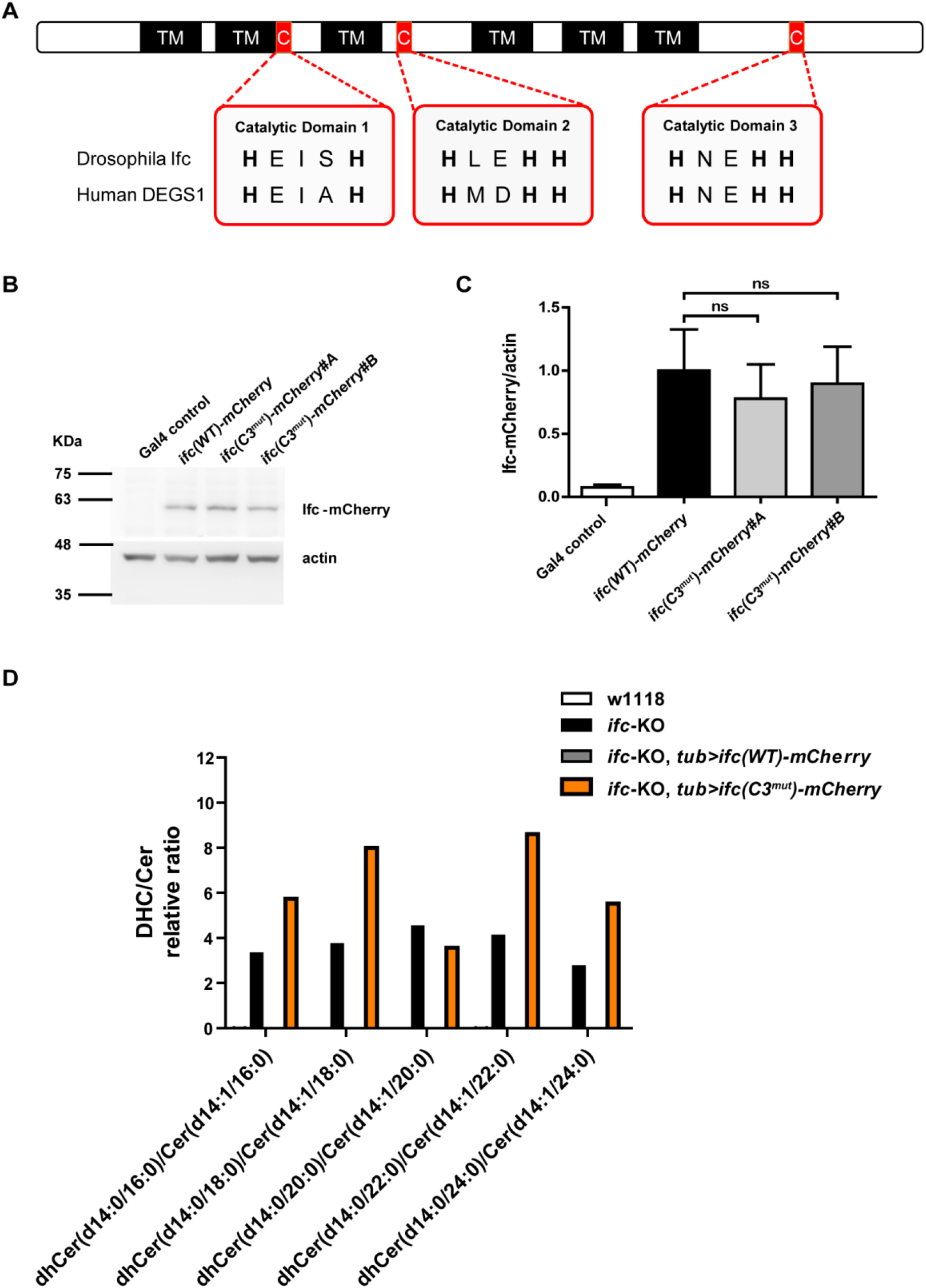
The construction and characterization of transgenically expressed *ifc(C3*^*mut*^*)-mCherry*. **(A)** A schematic illustration of the histidine-rich regions that are key catalytic domains of dihydroceramide desaturases. The amino acid sequences of the catalytic domains are highly conserved between *Drosophila* Ifc and Human DEGS1. TM: predicted transmembrane domains; C: histidine-rich catalytic domains. **(B)** Western blot analysis of Ifc-mCherry proteins expressed in flies under *nSyb-Gal4*. Actin was used as the loading control. **(C)** Quantification of the protein levels of mCherry-tagged *ifc* transgenes relative to actin. Normalized to *ifc(WT)-mCherry*. ns: Not significant (Student’s *t*-test). **(D)** The dhCer-to-Cer ratio of *w1118* control; *ifc-KO* alone; *ifc-KO,tub>ifc(WT)-mCherry*; and *ifc-KO,tub>ifc(C3mut)-mCherry* measured by HPLC-MS/MS.

**Figure S4.**
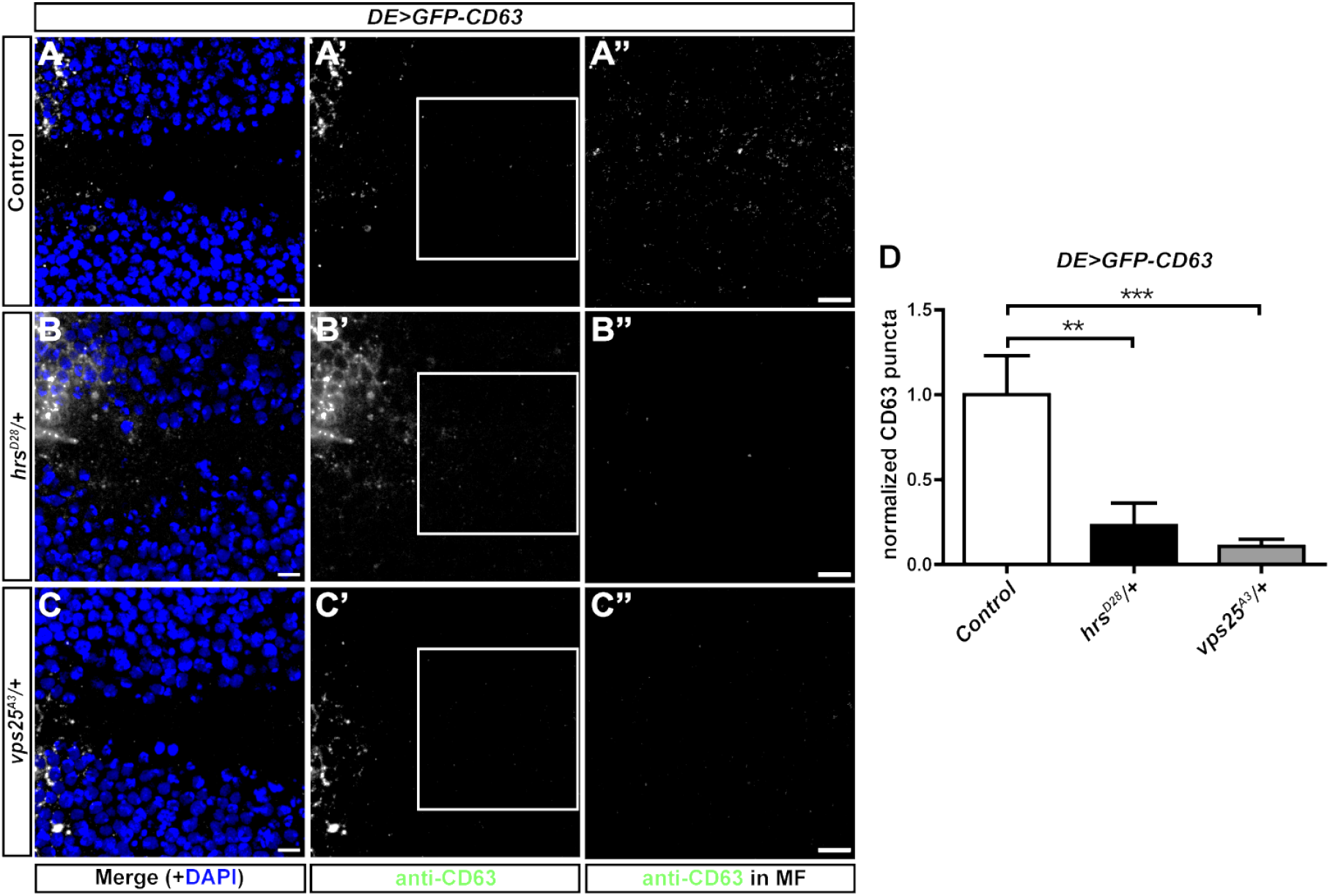
Heterozygous mutations in the ESCRT genes *hrs* and *vps25* reduce the number of exosomes. **(A-C)** Confocal images of *Drosophila* eye imaginal discs with dorsally expressed *GFP-CD63* in the wile-type control (A), *hrsD28/+* (B), and *vps25A3/+* (C) showing merged images with DAPI in blue (A,B,C), anti-CD63 staining in white (A’,B’,C’), and enlarged images of the ROI (white box) with linearly enhanced brightness and contrast to reveal GFP-CD63 puncta in the MF in the ROIs (A”,B”,C”). Scale bars: 10 μm. **(D)** Quantification of GFP-CD63 puncta. Bar charts show mean ± SEM of at least three independent experiments. *** p < 0.001; ** p < 0.01 (one-way ANOVA).

**Supplemental video 1**

Live imaging by spinning disc confocal microscopy of GFP-CD63 puncta produced by eye imaginal disc over-expressing Ifc(WT)-mCherry *ex vivo*.

**Supplemental video 2**

Live imaging by spinning disc confocal microscopy of GFP-CD63 puncta produced by mCherry-CAAX control eye imaginal disc.

**Supplemental video 3**

Light-sheet microscopy of GFP-CD63 puncta produced by eye imaginal disc over-expressing Ifc(WT)-mCherry.

**Supplemental video 4**

Light-sheet microscopy of GFP-CD63 puncta produced by the mCherry-CAAX control imaginal disc.

## Notes

### Competing Interest Statement

The authors have declared no competing interest.

### Summary of Updates

The revision adjusts the content to focuse on exosomes only.

